# Functional MRI of the Human Hippocampus at 10.5T: Pushing the Boundaries of Spatial Resolution

**DOI:** 10.64898/2026.04.30.721173

**Authors:** Yulia Lazarova, Lasse Knudsen, Steen Moeller, Nils Nothnagel, Alireza Sadeghi-Tarakameh, Lonike K. Faes, Noam Harel, Essa Yacoub, Kamil Uğurbil, Luca Vizioli

## Abstract

Although the hippocampus is central to human cognition, its deep location within the medial temporal lobe has historically limited fMRI signal quality. Here, we leverage an ultrahigh-field 10.5T fMRI system, which provides inherently higher signal-to-noise ratio than commonly used lower-field systems, to overcome these limitations and achieve unprecedented 0.5 mm isotropic resolution. This advancement allows for comprehensive hippocampal coverage with robust image quality, facilitating the study of functional microcircuits in individual subjects and paving the way for high-precision clinical diagnostics.

## Main

The hippocampus is key for human memory and cognition^1^ and supports diverse cognitive processes^2,3^, including declarative^4^, episodic^5^, and semantic^6^ memory, as well as statistical^7^ and associative learning^8,9^ and pattern completion^10^. These functions position the hippocampus at the apex of higher-order cognitive processes, that are particularly developed in humans. It comprises multiple subfields, including the subiculum, cornu ammonis (CA1, CA2, CA3/4), and dentate gyrus (DG)^11^, with distinct cytoarchitecture and specialized functional contributions^11–14,8^. As part of the allocortex, the hippocampus contains three histologically distinct layers, that differentially mediate information exchange with cortical and subcortical regions^11^.

Hippocampal dysfunction is implicated in multiple disorders and pathologies^15^ such as Alzheimer’s disease, age-related neurodegeneration, psychosis, depression, and epilepsy^16^. Understanding the mechanisms underlying these pathologies requires mesoscopic-scale study of hippocampal functional organization across subfields and individuals - a challenging feat even for more accessible cortical structures and resolutions. Such insight would enable interrogation of the specialized contributions of individual subfields to complex processes^8,17,18^, and holds transformative potential for both basic and translational neuroscience, yet remains poorly understood due to several methodological and technological challenges.

This gap partially reflects signal-to-noise ratio (SNR) limitations of functional Magnetic Resonance Imaging (fMRI) – the most widely used noninvasive brain imaging technique^19^– which directly constrains functional sensitivity, particularly when imaging subcortical structures that are away from the receive coils^20^, such as the hippocampus. Additional challenges arise from the convoluted hippocampal geometry and its thin gray matter ribbon, measuring approximately 2 mm in some subregions^21^, along with subfield-defining anatomical landmarks less than 1 mm thick^12^. These features require high spatial resolution which further reduces SNR. These challenges call for an fMRI approach that provides sub-millimeter resolution while maintaining high SNR. Ultra-high field (UHF; ≥7T) imaging offers a potential solution by providing supralinear SNR gains with increasing field strength, enabling submillimeter acquisitions^22–24^. Such resolution promises non-invasive investigations of mesoscopic functional organizations underlying fundamental units of neural computations across hippocampal subfields.

Another major obstacle to advancing our understanding of the hippocampus and its individual subfields is the lack of a universally accepted protocol for hippocampal subfield segmentation^25^ (for recent advances see ^26,27^), complicating cross-study comparisons and contributing to inconsistent findings^28,29^. Existing standardized protocols rely exclusively on anatomical images, because in conventional echo-planar imaging (EPI), the high spatial resolution needed to resolve hippocampal subfields entails a trade-off with functional contrast-to-noise ratio (fCNR) and spatial fidelity. However, cross-modal alignment is error-prone, particularly in submillimeter studies and in deep brain regions such as the hippocampus. Direct segmentation in functional space would also avoid inconsistencies arising from scan-dependent variability in cortical thickness and across acquisitions and sequences^30^. Developing EPI acquisitions with sufficient structural contrast to enable segmentation directly in functional space therefore represents a critical methodological goal for hippocampal fMRI.

Despite the strong biological motivation, submillimeter fMRI studies of the human hippocampus remain scarce. This may partially reflect the sparse and spatially distributed coding typical for hippocampal place cells^31^ for example, whereby only a small fraction of neurons are active at a given time without strong anatomical clustering, potentially producing weak and heterogeneous blood-oxygen-level-dependent (BOLD) signals^32^ that are difficult to detect at conventional resolutions due to partial volume effects. Existing UHF studies have predominantly focused on coarse anatomical divisions, such as anterior-posterior hippocampus^33,34^. In the few studies examining laminar effects, emphasis has primarily been on hippocampal vasculature ^35,36^, and physiological characterization using proof-of-concept tasks, rather than hypothesis-driven neuroscientific questions. This scarcity highlights the substantial technical challenges of in-vivo hippocampal imaging and the need to push spatial resolution beyond the current state-of-the-art (∼0.8mm isotropic^17,35–37^).

Here, we combined a unique 10.5T scanner, a custom-built 16-channel transmit/80-channel receive head coil^38^, tailored reconstruction techniques, and advanced denoising^39^ to leverage the higher (temporal) SNR and fCNR. We acquired functional BOLD images at an unprecedented 0.5 mm isotropic resolution from deep subcortical regions during visual stimulation in four individual participants. Our results demonstrate the feasibility of reliable single-subject functional mapping, enabling EPI-based subfield delineation without anatomical scans and highlight the potential of field strengths beyond 7T to advance human hippocampal fMRI.

### Results

By leveraging the supralinear gains in SNR and fCNR with increasing field strength^20,40^ we acquired whole-hippocampus EPI at 0.5mm isotropic resolution at 10.5T, achieving a fourfold reduction in voxel volume (0.125 µL) relative to the highest functional resolution reported previously (0.512 µL)^35,36^. Our 3D Gradient-Echo (GE) EPI protocol acquired 40 slices (Fig. 1a) (see <u>Online Methods</u>), providing full coverage of the hippocampus and parts of the visual, entorhinal and parahippocampal cortices. We employed a block design with 24-second ON and OFF intervals to present participants with scenes containing embedded objects that were either semantically congruent or incongruent with the background (Extended Data Fig. 1). We recorded 50 minutes of functional scanning for each participant, yielding sufficient data to generate high spatial fidelity maps that support single-subject analyses of hippocampal functional organization.

**Fig. 1.**
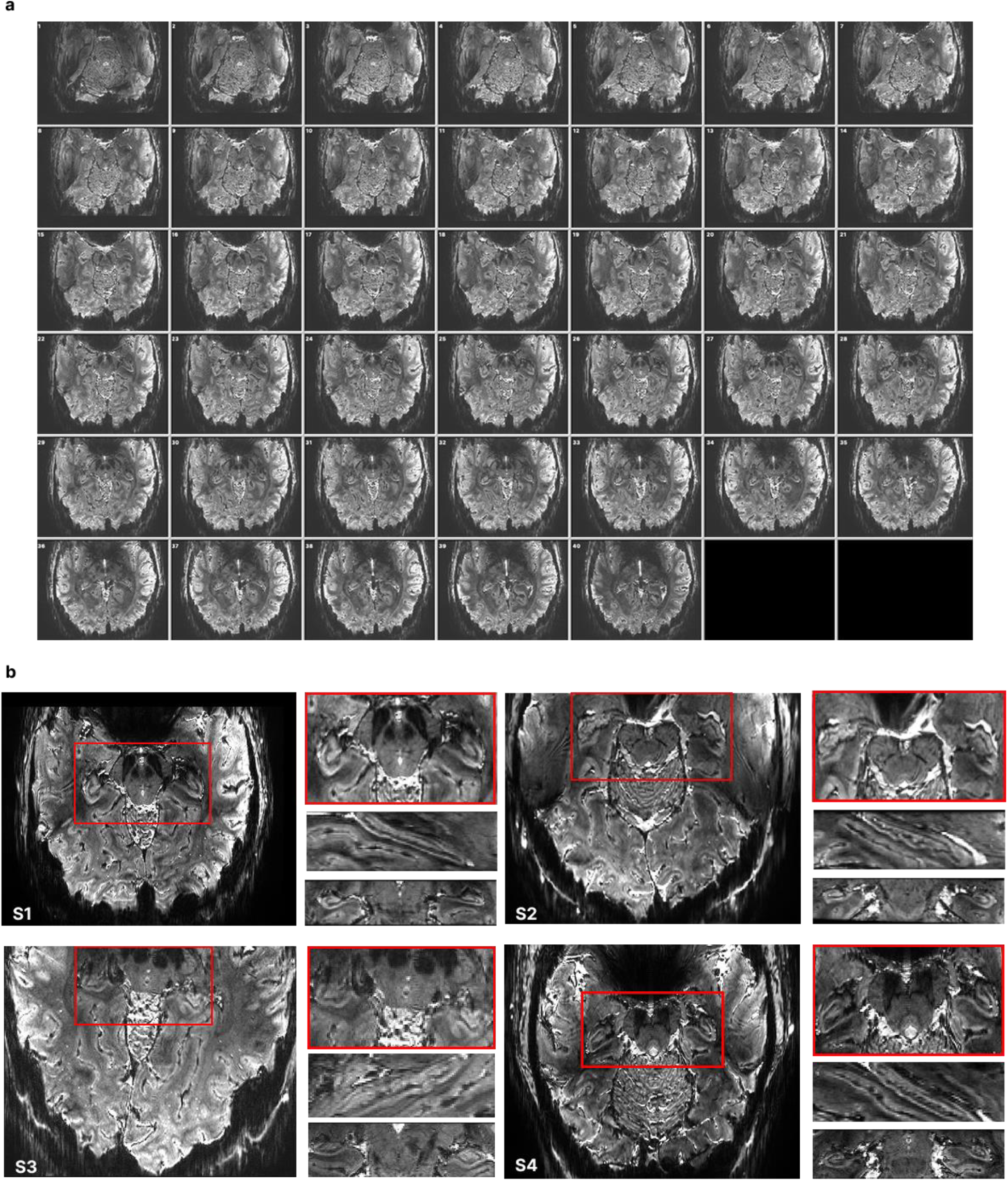
EPI at 0.5mm^3^ resolution. **(a)** EPI slices for one participant, illustrating the extent of the coverage achieved by the 40 slices: starting from slice 1 in the top left corner – most inferior point of our coverage, up to slice 44 in the bottom right corner – the most superior point of the brain covered. **(b)** Single-run mean EPI zoomed in over the hippocampus - averaged across a single run for each participant.

We observed clear structural image contrast (**Fig. 1**) in our EPI data, despite the unparalleled spatial resolution achieved. Manual subfield and laminar segmentation was performed directly in functional space based on single-run mean EPI data (**Fig. 1**b), enabled by the high structural detail of our functional images (**Fig. 2**a). We identified the subiculum, the DG and CA1 - CA3 subfields and performed layer segmentation for each participant (**Fig. 2** b, c).

**Fig. 2.**
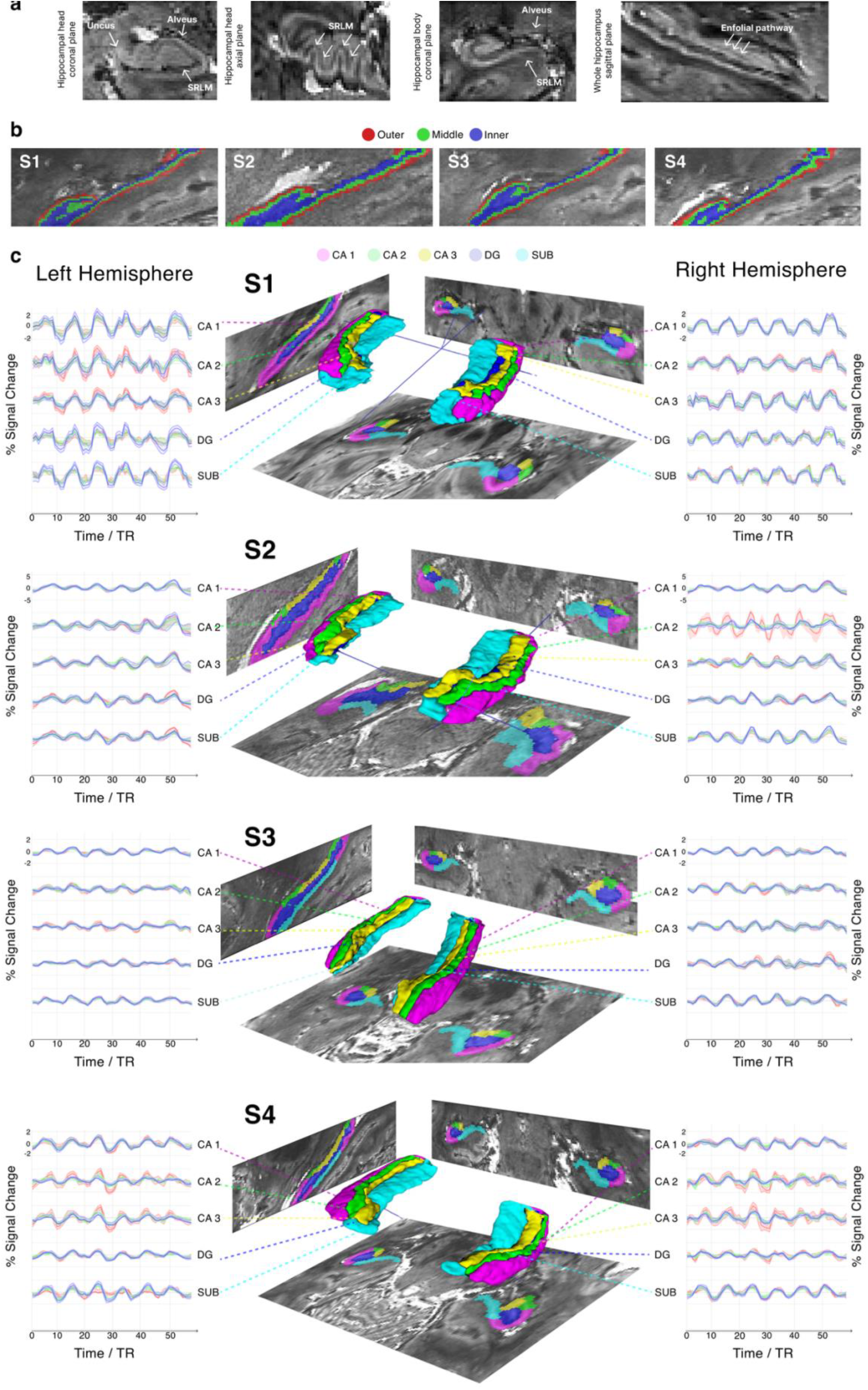
Hippocampal subfield segmentation and layer-specific time courses. **(a) Landmarks visible on mean-EPI**. Coronal and axial cut through the hippocampal head (first and second panel) and coronal cut through the body (third panel), demonstrating sufficient structural detail to locate some of the landmarks used to guide hippocampal subfield segmentation. The arrows indicate the location of the Alveus, Uncus, the Stratum Radiatum Lacunosum Moleculare (SRLM), and the endfolial pathway (longitudinal view – fourth panel). (b) **Laminar segmentation** of the hippocampus for each participant into three depths: inner (blue), middle (green) and outer(red). **(c) Hippocampal subfield-segmentations, 3D reconstructions and corresponding time courses across layers**. Five hippocampal subfields (DG– blue; CA1–magenta; CA2–green; CA3–yellow; subiculum (SUB) - cyan) superimposed over EPI data for each participant (middle column) with a 3D reconstruction. Columns 1 and 3 show the layer-specific time courses per subfield and hemisphere (left hemisphere – column 1; right hemisphere – column 3). The solid line shows the 20% trimmed mean percent signal change over time averaged across 10 runs; the shaded areas indicate standard error across runs. The correspondence to the layers is color-coded as seen in panel b). Refer to Table 1 in Extended Data for reference to the number of voxels contained in each subfield per layer.

We show subject-specific depth-resolved BOLD time courses, exhibiting the expected temporal characteristics, peaking during task blocks and returning to baseline during rest (**Fig. 2**c) for all subfields across hemispheres. **Fig. 2**c shows mean time courses calculated across 50 minutes of scanning for each participant, with shaded areas including standard error across runs.

We consistently achieved high temporal SNR (tSNR) in the hippocampus across participants (**Fig. *3***a). The mean voxel-wise tSNR values for an exemplary run from each participant indicate generally high values: (Mean ± SD tSNR across voxels) S1 - 23.6±12.4; S2 - 23.5±13.4; S3 - 42.6±21.2; S4 - 34.8 ±17.5, consistent with stable signal quality.

**Fig. 3.**
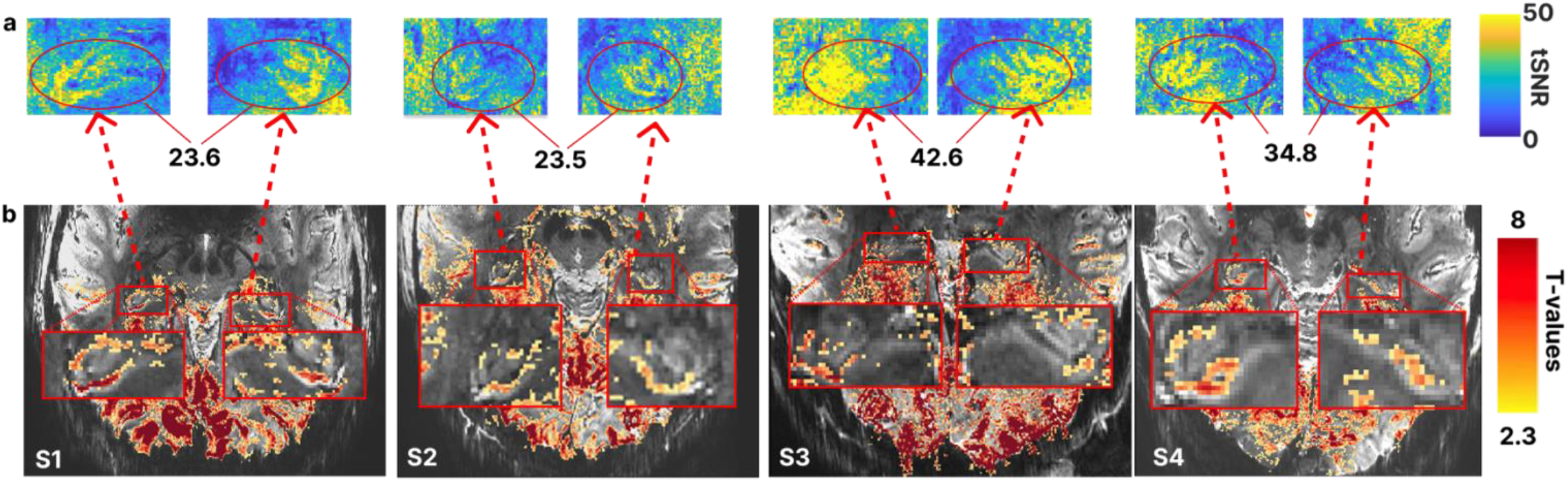
Temporal SNR and activity (t-value) maps. **(a) Hippocampus** tSNR maps for a single exemplar run per participant after NORDIC denoising, motion correction and linear detrending. The color indicates the mean tSNR across voxels for the whole hippocampus, averaged across both hemispheres. **(b)** EPI for each participant, overlayed with activity map from 10 runs showing positive responses to stimuli>baseline (FDR corrected, q<0.05). For visualization purposes only, activation maps were smoothed with a 1-voxel full width at half maximum (FWHM) kernel.

Finally, we observed spatially salient (i.e. following the hippocampal gray matter ribbon) and statistically significant (corrected for multiple comparisons using a false discovery rate (FDR) q<0.05) BOLD activations on subject-specific level for approximately 50 minutes of data (S1: t(577)=2.6, p=0.009; S2: t(577)=2.3, p=0.022; S3: t(577)=2.3, p=0.017; S4: t(577)=2.4, p=0.015). **Fig. *3***b shows the activation maps across participants, thresholded at t(577)=2.3 for consistency across subjects.

## Discussion

Our findings have profound implications for studying human hippocampus noninvasively. The advances achieved by our approach permitted reliable functional mapping of the hippocampus with unprecedented spatial resolution. Crucially, the robust and replicable high-quality functional maps and stable time courses permit analyses at the individual subject level and without relying on anatomical scans for subfield delineation. This is an elusive, yet crucial feat, even at standard resolutions.

The structural detail visible in our functional images approaches what has previously been achieved only with high-resolution anatomical scans^41^, enabling hippocampal subfield segmentation directly in functional space. This approach eliminates the need for cross-modal alignment between functional and anatomical data, an error-prone process that can critically compromise results especially in submillimeter studies. Defining depth-dependent regions of interest directly on the functional data further avoids inconsistencies arising from scan-dependent variability in cortical thickness, which can differ across acquisitions and choice of sequence^30^.

The 10.5T magnetic field strength affords high spatial resolution, enabling depth-resolved measurements in thin hippocampal structures. In regions where the hippocampal ribbon measures approximately 2 mm or less^21^, our 0.5mm isotropic resolution substantially increases the number of cortical depths samples compared to lower-resolution approaches. This enhances sensitivity to fine-grained functional variations, that may otherwise be obscured, particularly in the presence of spatial blurring ^42^ and partial volume effects.

Our approach, involving 10.5T and tailored reconstruction and NORDIC denoising^39^, achieved high tSNR values consistent across participants (**Fig. 2**a). These values surpassed the highest tSNR value reported in the literature (tSNR=21.84)^43^, which used substantially courser spatial resolution than that achieved in the present work (0.125 µL versus 0.512 µL voxel volume) and was optimized only for one hemisphere. Even at this enhanced spatial resolution, we observed robust, statistically significant and spatially coherent hippocampal activations at the single-subject level (**Fig. *3***b). Furthermore, extracted depth-resolved time courses from individual subjects (**Fig. 2**c) consistently revealed clear task-evoked responses across participants. This permits modelling of individual task epochs and supports reliable statistical inference at the single subject level by leveraging repeated measures across trials.

Acquiring high-quality, hippocampus-focused EPI data at this resolution helps alleviate a persistent limitation of the current literature: fMRI studies of the human hippocampus rarely display representative functional images, or voxel-wise activation maps, presumably due to low SNR in the hippocampal region. In contrast, we report single-subject activation maps in native functional space that display spatially salient and neuroanatomically plausible BOLD responses.

The combination of submillimeter spatial resolution, high SNR, and extended coverage achieved here represents an important step toward bridging microscale circuit organization and system-level imaging in the human brain. The acquired volume includes the full hippocampus along with portions of the entorhinal cortex and visual cortices. This extensive coverage, combined with unprecedented resolution for this cortical region, and a high level of (t)SNR permit concurrent assessment of depth-resolved hippocampal signals alongside cortical network activity. Invasive and histological studies have established that distinct laminar pathways within the hippocampus support different computational functions and connectivity patterns^44,45^. The spatial resolution achieved here therefore creates new opportunities to investigate these depth-dependent processes noninvasively in the human brain. By enabling depth-resolved hippocampal measurements together with network-level coverage, the present results help address longstanding trade-offs hippocampal fMRI between spatial resolution, spatial stability, and brain coverage. This capability opens the door to closer integration between human fMRI, computational models^46,47^, and circuit-level findings from animal studies^48^.

Depth-resolved hippocampal fMRI could open new avenues for investigating the neural mechanisms underlying age-related memory decline and associated clinical pathologies. By enabling interrogation of activity associated with feedforward and feedback pathways within cortico-hippocampal circuits, such measurements could, in future work, support investigations of predictive processing mechanisms implicated in neuropsychiatric and neurological disorders^49^. This capability may help identify which components of hippocampal circuitry are selectively affected in disease. Noninvasive, depth-resolved assessment of hippocampal subfield function at the individual level could therefore provide more precise mechanistic insight and ultimately support the development of patient-specific diagnostic or therapeutic strategies.

## Conclusion

Here we demonstrate a practical solution to several longstanding challenges in human hippocampal fMRI. By leveraging ultra-high magnetic field strength (10.5T), together with custom-built coil and advanced reconstruction and denoising, we achieve precise, noninvasive characterization of hippocampal microcircuitry and its interactions with cortical networks, bringing human fMRI closer to resolving circuit-level organization in vivo. The reliability and specificity of these measurements create new opportunities for translational research, including subject-specific mapping of functional alterations, disease progression monitoring, and targeted interventions. These advances provide a foundation for future studies of healthy hippocampal function and its disruption across aging and disease, representing an important step toward noninvasive, circuit-level imaging of the human hippocampus at the level of individual subjects.

## Online Methods

### Participants

Four healthy volunteers (4 males, with a mean age of 41.5, SD = 9.88) participated in the study. They all were right-handed and had normal or corrected-to normal vision. The volunteers were compensated for their participation at $50 per hour. Written informed consent was obtained in accordance with a protocol approved by the University of Minnesota Institutional Review Board. The study adhered to all relevant ethical regulations for research with human participants.

### Stimuli, study design and procedure

The stimuli consisted of 120 color images of sport scenes from four categories: football pitches (soccer fields), basketball courts, tennis courts, and baseball fields. Each scene was paired with an object and was scaled to allow for the object to be positioned centrally and sit naturally in the scene. The size of scenes and object across stimuli was kept consistent across stimuli. The background scenes were selected to contain enough category-related contextual information. Objects were either contextually congruent (e.g., a football in a football field), incongruent (e.g., a ball of yearn in a football field), or semi-congruent – where sports objects were paired with mismatching scene (e.g., a football in a basketball court).

Stimuli were presented in a block design with 24-second ON-OFF intervals. During each stimulation block, 12 randomly selected images from the same category (congruent, semi-congruent or incongruent) were displayed at 0.5Hz. Participants completed 10 runs, each lasting 5.2 minutes and containing two stimulation blocks per condition in randomized order. Each run began and ended with a 24-sec long inter-trial interval consisting of a grey screen with a fixation cross. The central object was consistently presented in the same portion of the visual field, spanning approximately 5.6° of visual angle. A red fixation cross was positioned at the center of the screen, and participants were instructed to maintain fixation throughout the experiment.

Stimuli appeared centrally on the screen at a visual angle of approximately 5.6° and were presented using MATLAB (version R2021b) with Psychophysics Toolbox (version 3.0.15)^50^. Images were projected in the scanner using a VPixx PROPixx MRI projector (1440 Hz) connected to a Mac Pro (1920 × 1080 resolution). Participants viewed the screen through a mirror mounted in the head coil at distance of ∼89.5 cm. Behavioral responses were given using a right-hand MR-compatible VPixx 4-button box. Participants rated their level of surprise for each stimulus on a scale from 1 (not at all surprising) to 4 (very surprising). They were instructed to minimize any head movements throughout the experiment.

### Data acquisition

Functional MRI data were collected using a 10.5 T Siemens Magnetom System (Siemens Healthineers, Erlangen, Germany) with an custom-built 16-channel transmit and 80-channel receive head coil^38^. We used a 3D GRE EPI sequence, with a nominal resolution of 0.5mm isotropic voxels (0.125 µL); in-plane acceleration factor along the phase encoding direction (iPAT) factor of 4; repetition time (TR) 120ms; 40 slices; phase encoding direction: P>>A; volume acquisition time (VAT) of 5280ms; echo time (TE) of 26.2ms, partial Fourier (pF) 5/8, readout bandwidth of 850Hz/Px, and nominal Flip Angle (FA) of 20°. For each participant we performed manual shimming to improve B0 homogeneity over the hippocampal structure bilaterally.

### Tailored offline image reconstruction

All EPI data where processed with a three step phase correction: even/odd readout correction using a linear time correction estimated from a 3-line navigator using opposite polarity^51^, correction for apparent off-resonance motion from B0 (DORK^52^) estimated from the 3-line navigator using same polarity, and correction of apparent shot to shot bulk phase variations using a data self-consistency technique^53,54^. In-plane under-sampling was corrected using GRAPPA for k-space interpolation, and for phase-coherent channel combination a SENSE1 combination was used with sensitivity profiles obtained using ESPIRIT. Thermal noise was reduced using the NORDIC^39^ approach on the complex data and processed on each run independently. Magnitude data was used for further processing.

### Data preprocessing

All functional runs were motion-corrected in two steps: within-run alignment to the first volume of the run, and across-run alignment to run 6 (recorded approximately mid-way through the session, therefore serving as a mid-point between the head position at the start and at the end of the scanning session). The within-run alignment was performed in AFNI^55^ (*3dvolreg* function), using rigid-body transformation with three translation and three rotation parameters. Across-run alignment was performed in Advanced Normalization Tools (ANTs)^56^, where the mean of each run was calculated and aligned to the mean of the target run (run 6), using the *antsRegistration* function, which included both linear and non-liner transformations. Motion parameter estimation in both steps was restricted to a manually defined brain mask in ITK-SNAP^57^ (www.itksnap.org) to include only brain tissue. The resulting transforms from the two steps were concatenated and applied in one interpolation using the *3dNwarpApply* function in AFNI.

The first two volumes of each run were discarded in BrainVoyager^58^ to remove signal instability before steady-state was reached. Temporal high-pass filtering was then applied to remove low drifts using a general linear model (GLM) with a design matrix including up to the 3^rd^-order discrete cosine transform (DCT) basis set of predictors. Temporal smoothing was performed using ANTs’ *3dTsmooth* function with a three-point Hamming window.

No spatial smoothing was applied to preserve the high-resolution the fine-scale structural details. All subsequent analysis were conducted in functional space, therefore, no distortion correction or alignment to anatomical images was performed.

### Segmentation and ROI definition

Segmentation of hippocampal subfields was performed in ITK-SNAP directly on individual subjects’ functional echo-planar images, using a mean image from the motion correction target run (run 6).

We followed the segmentation procedures outlined in ^59^ and ^60^ – the only high-resolution protocols we found covering the entire hippocampus. We used these protocols in conjunction with ^60^ – a protocol based on 7T MRI images. Five subfields were defined: subiculum, cornu ammonis 1-3 (CA1, CA2, CA3), and dentate gyrus (DG). Additional guidance from ex vivo literature^61^ was used for decisions regarding points were protocols often diverge: (1) borders between CA1 and subiculum were defined following the procedure described in ^59^ and ^60^, minimizing the risk of mislabeling subiculum voxels as CA1; (2) voxels belonging to the stratum radiatum-lacunosum-moleculare (SRLM) were used to delineate the boundary between CA and DG, without assigning them a discrete label; where the SRLM exceeded 1 voxel in thickness, these voxels were assigned to the superiorly located structure^60^; (3) borders around CA2 and CA3 were guided by slices where the endfolial pathway was visible **Fig. 2a**), in conjunction with the geometrical principle described in ^60^ for areas lacking visible landmarks. The subiculum was defined as a single structure, without further subdivision.

### Layer segmentation

We defined three hippocampal layers, motivated by the three distinct layers of cells known in the allocortex^12^. Importantly, these layers represent depth-wise divisions derived from the imaging data and are not intended to imply a one-to-one correspondence with histologically defined cellular layers.

First, for each subregion we manually selected the outer and the inner border of the hippocampus directly onto the EPI data. We used the resulting file to execute LayNii’s ^62^ function LN2_LAYERS. For this purpose, the data was up-sampled to 0.25mm^3^ resolution, and subsequently resampled back into its original space after the layer definition. We used an equidistant approach for layer estimation, yielding three relative depth bins (inner, middle and outer) spanning the hippocampal thickness. The resulting file containing the three labels was intersected with the subregions defined in the previous step to create layers per subregion (refer to Extended Data Table 1 for information on the size of each layer per subregion).

### Data analyses

All analysis were performed at the single-subject level. In BrainVoyager (version 21.4) we computed stimulus-evoked functional maps across the entire imaging slab. For each participant maps were generated from all 10 runs, contrasting stimuli > baseline, and corrected for multiple comparisons using a false discovery rate (FDR) approach^63^ with a threshold q<0.05. Stimulus-evoked activations were quantified using a standard general linear model (GLM). The BOLD signal was modelled with a double-gamma hemodynamic response function and a ‘boxcar’ predictor corresponding to the stimulus onset and offset times.

Subsequent analyses were performed using custom MATLAB scripts (ver. R2018b, *Natick, Massachusetts: The MathWorks Inc*.) in combination with the *NeuroElf toolbox (v10_5153)* and *BVQXtools (ver. 08b)*. Individual voxel time courses were extracted from the 20% most active voxels within each ROI and run, then averaged across runs (solid lines Fig 2c). 20% trimming of the mean across voxels was applied. The shaded red area in Fig 2c represents the standard deviation (SD) across runs. Temporal SNR values were calculated by dividing the mean time-courses (over time) by the SD, computed independently per each voxel, run and participant.

## Acknowledgements

The authors thank Alexander Bratch and Matt Waks for the technical support throughout our data collection; and T.J. Whitney for her help with participant recruitment. This work was supported by NIH grants S10 RR029672, P41 EB027061, R01 EB038654, R01 NS136490 and P30 NS076408.

