## Supplementary Material for "Functional MRI of the Human Hippocampus at 10.5T: Pushing the Boundaries of Spatial Resolution"

### Extended Data

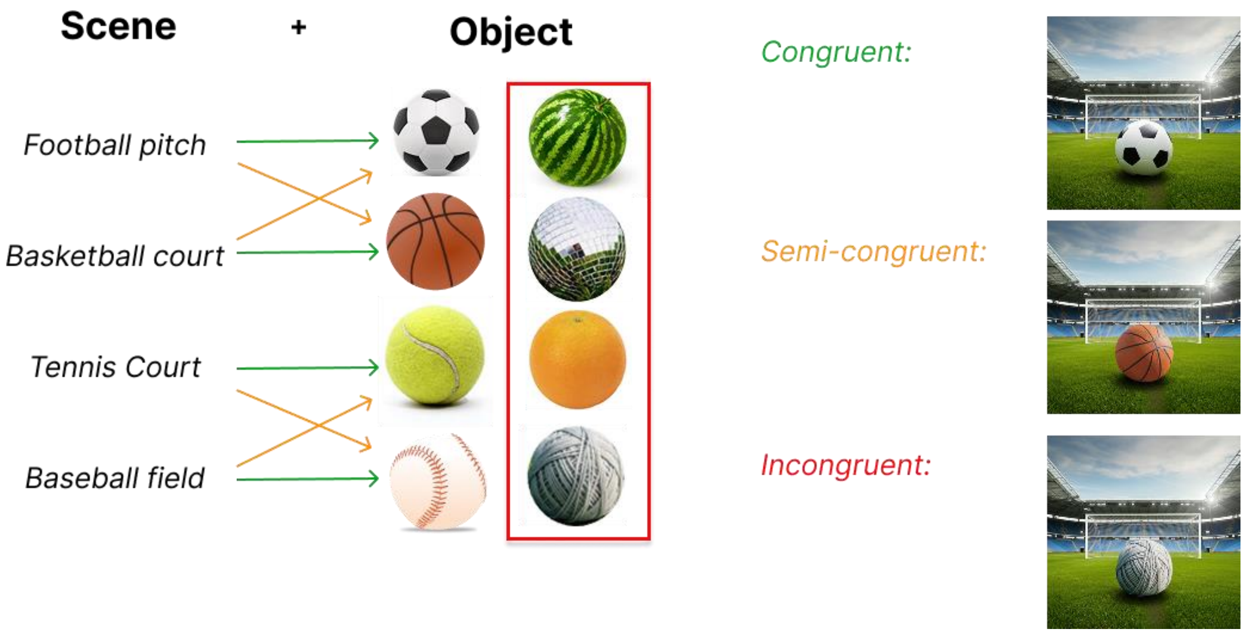


***Extended Data Fig. 1 Experimental paradigm and scene – object congruency conditions***
The study employed a block-design paradigm with 24 s stimulus blocks interleaved with 24 s baseline periods. Each congruency condition (congruent, semi-congruent, incongruent) was presented in a randomized order across 10 runs, with two repetitions per run, yielding 20 total presentations per condition. Stimuli were constructed by semi-randomly pairing 120 background scenes per category with focal objects. Objects were always positioned centrally at a fixed viewing angle to ensure natural embedding within the scene. To maintain size regularity and prevent preattentive size-based confounds, semi-congruent stimuli were restricted to sport-specific objects (e.g., basketball for football pitch scenes; baseball for tennis court scenes). Incongruent stimuli paired scenes with objects of similar physical dimensions but unrelated function (e.g., disco ball and watermelon for football pitch; orange and yarn ball for tennis court and baseball scenes). Green, orange, and red arrows indicate congruent, semi-congruent, and incongruent pairings, respectively. The red box highlights the incongruent object set. Representative example stimuli for each condition are shown in the rightmost column. All scene–object combinations were counterbalanced across runs to control for low-level visual properties and scene familiarity.

***Extended Data Table 1: Number of voxels across the three layers of all hippocampal subregions for each hemisphere of each participant.***

| Sub ID | ROI | Hemisphere | Layer | Nr. voxels |
|---|---|---|---|---|
| **S1** | CA1 | LH | Outer | 2598 |
| | | | Middle | 1553 |
| | | | Inner | 1234 |
| | | R H | Outer | 2634 |
| | | | Middle | 1826 |
| | | | Inner | 1697 |
| | CA2 | LH | Outer | 628 |
| | | | Middle | 542 |
| | | | Inner | 648 |
| | | RH | Outer | 571 |
| | | | Middle | 550 |
| | | | Inner | 749 |
| | CA3 | LH | Outer | 1336 |
| | | | Middle | 894 |
| | | | Inner | 852 |
| | | RH | Outer | 1012 |
| | | | Middle | 675 |
| | | | Inner | 553 |
| | DG | LH | Outer | 510 |
| | | | Middle | 1534 |
| | | | Inner | 3245 |
| | | RH | Outer | 312 |
| | | | Middle | 1150 |
| | | | Inner | 4147 |
| | Sub | LH | Outer | 2759 |
| | | | Middle | 2442 |
| | | | Inner | 2635 |
| | | RH | Outer | 2050 |
| | | | Middle | 1596 |
| | | | Inner | 1713 |
| **S2** | CA1 | LH | Outer | 2739 |
| | | | Middle | 1881 |
| | | | Inner | 1233 |
| | | RH | Outer | 3446 |
| | | | Middle | 2567 |
| | | | Inner | 1934 |
| | CA2 | LH | Outer | 791 |
| | | | Middle | 600 |
| | | | Inner | 332 |
| | | RH | Outer | 1048 |
| | | | Middle | 972 |
| | | | Inner | 809 |

| | | | | |
|---|---|---|---|---|
| | CA3 | LH | Outer | 1037 |
| | | | Middle | 1039 |
| | | | Inner | 523 |
| | | RH | Outer | 1009 |
| | | | Middle | 996 |
| | | | Inner | 698 |
| | DG | LH | Outer | 742 |
| | | | Middle | 1291 |
| | | | Inner | 2321 |
| | | RH | Outer | 1088 |
| | | | Middle | 1966 |
| | | | Inner | 3870 |
| | Sub | LH | Outer | 2909 |
| | | | Middle | 3218 |
| | | | Inner | 3653 |
| | | RH | Outer | 2264 |
| | | | Middle | 2609 |
| | | | Inner | 2697 |
| **S3** | CA1 | LH | Outer | 3125 |
| | | | Middle | 2407 |
| | | | Inner | 1862 |
| | | RH | Outer | 3330 |
| | | | Middle | 2616 |
| | | | Inner | 2153 |
| | CA2 | LH | Outer | 891 |
| | | | Middle | 711 |
| | | | Inner | 499 |
| | | RH | Outer | 895 |
| | | | Middle | 773 |
| | | | Inner | 690 |
| | CA3 | LH | Outer | 1356 |
| | | | Middle | 1318 |
| | | | Inner | 1040 |
| | | RH | Outer | 1243 |
| | | | Middle | 1117 |
| | | | Inner | 831 |
| | DG | LH | Outer | 603 |
| | | | Middle | 1214 |
| | | | Inner | 2087 |
| | | RH | Outer | 424 |
| | | | Middle | 983 |
| | | | Inner | 2273 |
| | Sub | LH | Outer | 2273 |
| | | | Middle | 2501 |
| | | | Inner | 2505 |

| | | | | |
|---|---|---|---|---|
| | | RH | Outer | 2011 |
| | | | Middle | 2386 |
| | | | Inner | 2330 |
| **S4** | CA1 | LH | Outer | 2185 |
| | | | Middle | 1713 |
| | | | Inner | 1271 |
| | | RH | Outer | 2434 |
| | | | Middle | 1893 |
| | | | Inner | 1295 |
| | CA2 | LH | Outer | 476 |
| | | | Middle | 497 |
| | | | Inner | 355 |
| | | RH | Outer | 565 |
| | | | Middle | 565 |
| | | | Inner | 505 |
| | CA3 | LH | Outer | 886 |
| | | | Middle | 871 |
| | | | Inner | 600 |
| | | RH | Outer | 720 |
| | | | Middle | 610 |
| | | | Inner | 428 |
| | DG | LH | Outer | 501 |
| | | | Middle | 1090 |
| | | | Inner | 2063 |
| | | RH | Outer | 320 |
| | | | Middle | 856 |
| | | | Inner | 2286 |
| | Sub | LH | Outer | 1449 |
| | | | Middle | 1721 |
| | | | Inner | 2256 |
| | | RH | Outer | 1964 |
| | | | Middle | 1994 |
| | | | Inner | 2016 |
